# *Nr5a1* suppression during the fetal period optimizes ovarian development by fine-tuning of Notch signaling

**DOI:** 10.1101/331090

**Authors:** Risa Nomura, Kenichi Kashimada, Hitomi Suzuki, Liang Zhao, Atsumi Hosokawa-Tsuji, Hideo Yagita, Masatoshi Takagi, Yoshiakira Kanai, Josephine Bowles, Peter Koopman, Masami Kanai-Azuma, Tomohiro Morio

## Abstract

The nuclear receptor NR5A1 (also known as Ad4BP, or SF1) is essential for the initial steps of mammalian gonadal development. The Nr5a1 gene is equally expressed in XX and XY gonadal primordia, but after sex determination, is up-regulated in XY and down-regulated in XX gonads. We recently reported a case of 46, XX disorder of sex development (DSD) in which ectopically expressed NR5A1 in XX gonads led to an ovo-testicular phenotype, suggesting that excess NR5A1 can direct the development of immature XX gonads towards testicular formation. However, a direct causal relationship has not been demonstrated in an animal model. Here, using a Wt1-BAC (bacterial artificial chromosome) transgene system, we generated two lines of mice overexpressing Nr5a1 in the fetal gonads at different levels. One of these lines (Tg-S), highly expressing Nr5a1, revealed that enforced Nr5a1 expression alone is insufficient to switch the fate of the 46,XX gonads toward testicular formation in mice. In the other line (Tg-A) expressing Nr5a1 at lower level, ovarian development was compromised, with multi-oocyte follicles, reduced number of matured follicles, and impaired expression of Wnt4, resulting in late onset infertility at 20 weeks after birth. The phenotype was similar to that of genetically modified mice with impaired Notch signaling. Indeed, the expression level of Notch2 and 3 was significantly reduced in Tg-A mice, and the ovarian phenotype in Tg-A mice was almost completely rescued by in utero treatment with a Notch2 agonist HMN2-29. We conclude that suppression of Nr5a1 during the fetal period optimizes ovarian development by fine tuning of Notch signaling levels.

**AUTHOR SUMMARY:** Sexual development is a process of differentiation from undifferentiated bipotential gonads, and insight into sexual differentiation will bring important new knowledge to our understanding of organogenesis. The nuclear receptor NR5A1 which is essential for mammalian gonadal development, is equally expressed in both gonadal primordia, but after sex determination, is up-regulated in XY and down-regulated in XX gonads. We have recently demonstrated that this down-regulation is mediated by ovarian transcription factor, Forkhead box L2 (FOXL2). This finding raised two key questions, whether Nr5a1 can function as a male sex-determining factor, and whether the repression is essential for appropriate ovarian development. By generating two lines Tg mice in XX gonads with different enforced expression levels of Nr5a1, our present study revealed that alterations in Nr5a1 dosage, either reduced or excessive, result in pathological effects in ovarian development and female fertility, indicating that the precise control of Nr5a1 at the transcriptional level is essential for optimal ovarian development. We envisage that the improved understanding of how this pathway regulates ovarian development and female fertility would aid the development of artificial somatic ovarian cells, which in turn may provide a valuable treatment option in reproductive medicine.

**ABBREVIATIONS:**

BAC: bacterial artificial chromosome
dpc: days post coitum
FOXL2: Forkhead box L2
HSD: hydroxysteroid dehydrogenase
IF: Immunofluorescence
MOFs: multiple oocyte follicles
PFA: paraformaldehyde
qRT-PCR: Quantitative real-time PCR
Rps29: Ribosomal protein S29
SRY: sex-determining region Y

## INTRODUCTION

NR5A1, also known as Ad4BP or SF1, is a member of the nuclear receptor superfamily. In mice, *Nr5a1* is expressed from about 9.5 days *post coitum* (dpc), in the anlagen of the gonads and the adreno-genital primordium [1], and knockout models show complete gonadal agenesis in both XX and XY, suggesting that NR5A1 is essential for genital ridge development in both sexes [2] [3]. In addition to initiating gonadal development, NR5A1 plays crucial roles in testicular development. It is transcriptionally up-regulated in the developing mouse testes [4], where it acts as a cofactor of the male sex-determining factor SRY (sex-determining region Y) to induce *Sox9* expression [5]. Subsequently, it cooperates with SOX9 to maintain the expression of *Sox9* itself [5] and up-regulate other Sertoli cell-specific genes, including *AMH* (De Santa Barbara et al., 1998). Furthermore, NR5A1 is essential for the differentiation of the testicular steroidogenic cells, fetal Leydig cells [6]. Consistent with its essential roles in testis differentiation, heterozygous loss-of-function mutations in *NR5A1* cause XY female sex reversal in humans [7-10].

In contrast to its transcriptional up-regulation in fetal testes, *Nr5a1* expression in mouse fetal ovaries decreases after 12.5 dpc [1]. We have recently demonstrated that this down-regulation is mediated by Forkhead box L2 (FOXL2), a key ovarian transcription factor expressed mainly in pregranulosa/granulosa cells [11]. FOXL2 directly binds to the proximal promoter of *Nr5a1*, thereby antagonizing the actions of WT1-KTS. Given the highly conserved sequence for FOXL2 binding, this regulatory mechanism may be maintained in other eutherian mammals [11]. Based on these observations, we hypothesized that adequate suppression of *Nr5a1* may be essential for normal ovarian development. However, the biological relevance of *Nr5a1* suppression during fetal ovarian development has not been clarified.

Recently, we and others have reported 46,XX testicular or ovo-testicular DSD individuals carrying mutations in codon 92 of *NR5A1*, including the R92W and R92Q mutations [12-16]. The variant protein was thought to function in the XX gonads by escaping the suppressive action of NR0B1 (DAX-1), a pro-ovary factor [13, 16, 17]. The presence of testicular tissue in the probands further suggested that ectopic activity of NR5A1 may drive testis differentiation in the absence of the *SRY* gene.

These observations raised two key questions. Firstly, it is not clear whether *Nr5a1* can function as a male sex-determining factor, i.e., whether elevated *Nr5a1* expression levels in XX gonads where the male determining gene *Sry* is absent, are sufficient to direct the fate of gonads towards testicular development. Secondly, it is not known whether repression of *Nr5a1* is essential for appropriate ovarian development.

To address these questions, we exploited a BAC (bacterial artificial chromosome) transgene system [18] whereby *Nr5a1* expression is driven by *Wt1* regulatory sequences. In the fetal XX gonads, endogenous *Wt1* is expressed in supporting cells including pre-granulosa cells and coelomic epithelium [19-21]. Hence, by directing transgenic *Nr5a1* expression to *Wt1*-expressing XX gonadal supporting cells, we aimed to investigate the consequence of *Nr5a1* overexpression in the relevant cell types in XX mouse fetal gonads. Molecular and phenotypic analysis of the two transgenic mouse lines generated demonstrated, firstly, that enforced *Nr5a1* expression alone is insufficient to switch the fate of the 46,XX gonads toward testicular formation in mice and, secondly, that overexpression of NR5A1 disrupts ovarian follicular development and causes premature ovarian insufficiency by dysregulating levels of Notch signaling, which is known to be important for ovarian development and function.

## RESULTS

### Overexpression of *Nr5a1* fails to cause XX sex reversal in mice

Using the piggyBac-enabled *Wt1*-BAC system [18] (Fig. 1A), we successfully generated two transgenic mouse lines (Tg-A and Tg-S) expressing an *Nr5a1-*IRES-*Egfp* transgene at different levels (Fig. 1B–M). Using quantitative reverse transcriptase PCR (qRT-PCR) we found that XX Tg-S fetal gonads expressed *Nr5a1* at almost the same level as that in wild type testes, whereas the XX Tg-A gonads expressed *Nr5a1* at levels between those in wild type testes and ovaries (Fig. 1N). We confirmed histologically that the kidneys and adrenal glands, expressing endogenous *Wt1* at high levels, developed no apparent abnormalities in either Tg-A or Tg-S mice, suggesting that the gonadal phenotypes of the transgenic mice were unlikely to be caused by the impaired function of those organs.

**Figure 1:**
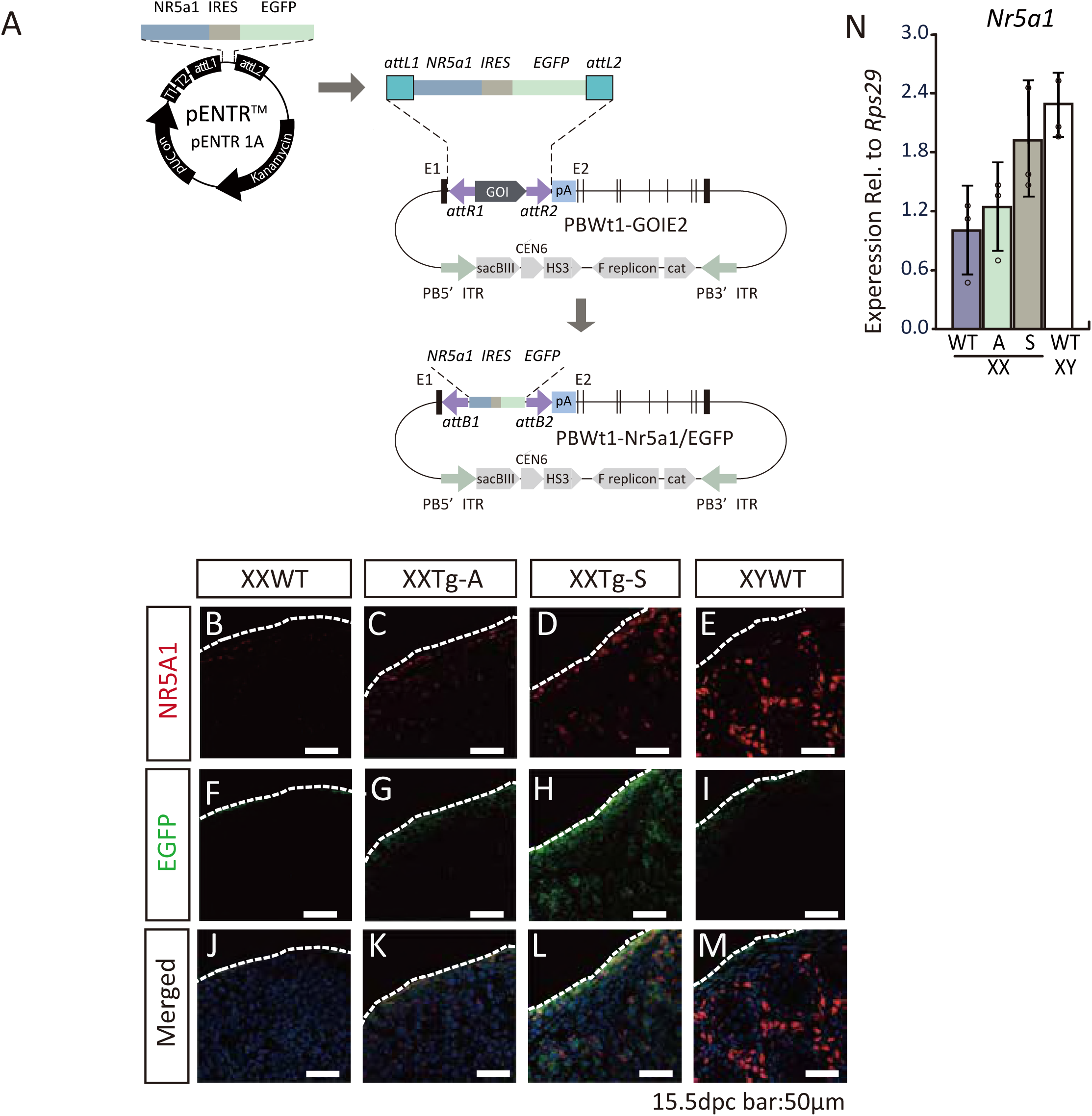
Generation of transgenic mice with enforced *Nr5a1* expression in XX gonads. (A) Schematic representation of the strategy to generate transgenic mice using the piggyBac-enabled *Wt1*-BAC system. *Nr5a1*-IRES-*Egfp* fragment in the Gateway entry vector was inserted into PBWt1-Dest using LR recombination. (B-M) Transgenic EGFP expression was detected using immunofluorescence in the XX fetal gonads at 15.5 dpc in both Tg-A and Tg-S lines. EGFP and NR5A1 were co-localized in XX transgenic gonads (K, L). (N) qRT-PCR analysis showing that *Nr5a1* was expressed at different levels in the XX fetal gonads of the Tg-A and Tg-S lines at P0. In the XX fetal gonads of the Tg-S line, *Nr5a1* was expressed at levels similar to that in wild-type testes. In the Tg-A line, *Nr5a1* was expressed in the XX fetal gonads at levels between those in wild-type testes and ovaries. Expression levels were normalized to *Rps29.* Mean ± SD, n = 3.

In both Tg-A and Tg-S lines, XX mice developed female internal and external genitalia (Fig. 2A–F). Morphologically, the gonads in adult XX Tg-S mice were streak-like and elongated (Fig. 2F,J). In Tg-A mice, fetal ovaries were longer and thinner than wild type (Fig. S1A-D), However, the elongation of the gonads resolved by P0. Since a similar phenotype was reported in *Sox4-/-*mice [22], we examined the expression levels of *Sox4* in the XX Tg-A gonads at 15.5 dpc and indeed found significantly reduced expression of *Sox4* (Fig. S1E), suggesting that elevated NR5A1 repressed *Sox4* resulting in abnormal morphology of the fetal ovaries.

**Figure 2:**
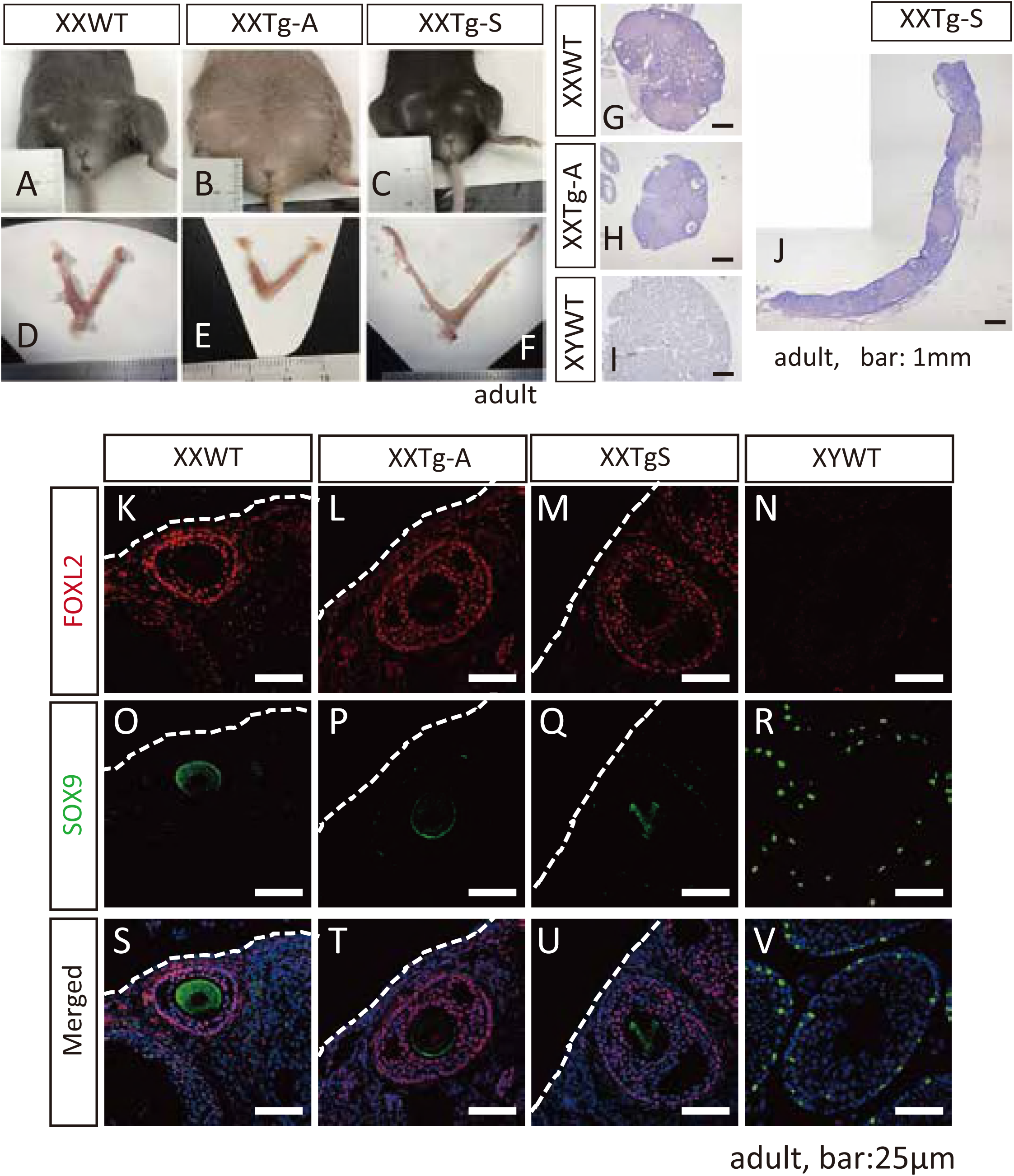
Transgenic overexpression of *Nr5a1* in XX fetal gonads did not induce male development. (A–C) External genitalia of wild-type, Tg-A and Tg-S XX mice at 6–8 months old. XX mice in both transgenic lines developed female-type external genitalia. (D–F) Reproductive tracts of wild-type, Tg-A, and Tg-S XX mice at 6–8 months old. In the XX Tg-S mice, the uterine horn and ovaries were elongated compared to wild type (F). (G–J) Histological analysis of gonadal sections at 6-8 months using HE stain. XX gonads of both transgenic lines contained ovarian follicles but no seminiferous tubule-like structures. (K–V) Immunofluorescence analysis for markers of granulosa (FOXL2, in red) or Sertoli cells (SOX9, in green) in the fetal gonads in adult mice. In the XX gonads of the Tg-A and Tg-S lines, only FOXL2-positive granulosa cells were present (L,M), and no SOX9-positive cells were detected (P, Q; some non-specific staining of the oocytes was seen using the anti-SOX9 antibody).

In both transgenic lines, histological examination of XX adult gonads revealed the presence of ovarian follicles but detected no seminiferous tubule-like structures (Fig. 2G-J). Consistently, FOXL2, a marker of ovarian granulosa cells, was expressed in XX adult gonads in both mouse lines (Fig. 2L, M). In contrast, SOX9, a marker of testicular Sertoli cells, was not detected (Fig. 2P, Q). Based on these data, we conclude that *Nr5a1* overexpression alone is insufficient to cause XX sex reversal in mice.

### Presence of ectopic steroidogenic cells in fetal ovaries of *Nr5a1*-trangenic mice

In addition to Sertoli cells, NR5A1 is highly expressed in gonadal steroidogenic cell lineages in both sexes and is essential for their differentiation (Buaas Swain, Development 2012) [23, 24]. In the males, steroidogenic Leydig cells differentiate during fetal testis development and produce androgen [25]. In contrast, the ovarian steroidogenic theca cells do not differentiate until after birth [25-27]. We asked whether overexpression of *Nr5a1* is able to induce the ectopic differentiation of steroidogenic cells in fetal XX gonads. Immunofluorescence analysis of the XX fetal gonads of Tg-A and Tg-S mice (Fig. 3A–L) revealed that 3β-hydroxysteroid dehydrogenase (3β-HSD), the enzyme mediating the first step of steroidogenesis and a marker of steroidogenic cells [28], was detected in the Tg-S XX gonads (Fig 3G). However, the plasma testosterone and estradiol levels at P0 were not elevated in XX Tg-S mice compared to wild type (Fig. 3M, N). This is consistent with the lack of masculinization in XX Tg-S mice (Fig. 2C,F).

**Figure 3:**
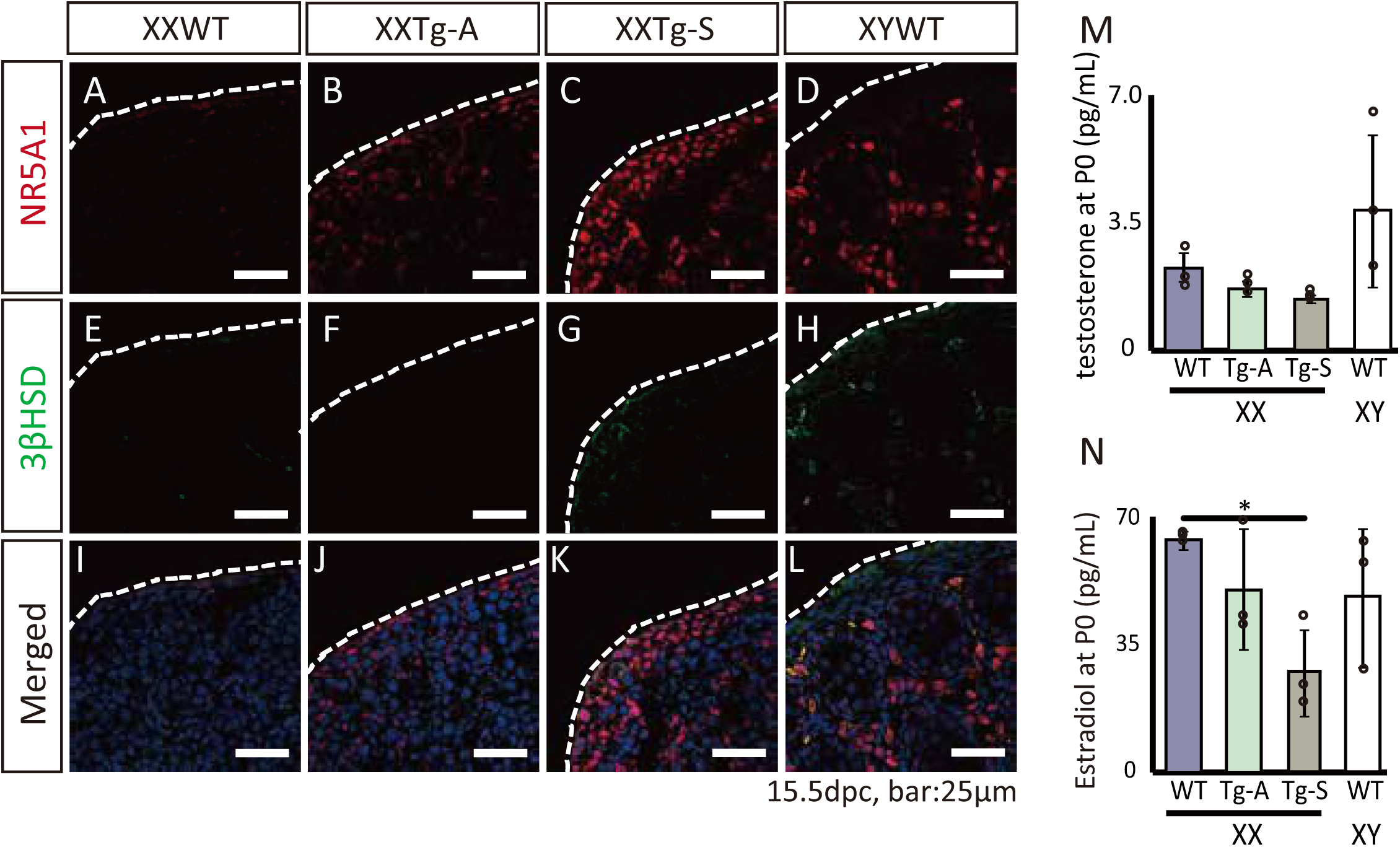
Formation of ectopic steroidogenic cells in the XX transgenic fetal gonads. (A–L) Immunofluorescence analysis for 3β-HSD at 15.5dpc, a marker of steroidogenic cells. In the Tg-S fetal ovaries, 3β-HSD positive cells were sparsely observed, some of which showed co-localization with NR5A1 (G,K). (M, N) The plasma testosterone (M) or estradiol (N) levels at P0. No significant increase in levels of testosterone or estradiol was observed in XX Tg-A or Tg-S mice compared with wild-type female mice. Mean ± SEM, n = 3. \**p* < 0.05 (Welch‘s *t*-test).

### Enforced expression of *Nr5a1* leads to increased follicular atresia and impaired fertility

Histological examination of Tg-A and Tg-S ovaries revealed increased number of multiple oocyte follicles (MOFs) (Fig. 4A–D), suggesting a disruption of ovigerous cord fragmentation. Further, we found that the numbers of antral follicles (type 6–8) decreased significantly in both transgenic lines compared to wild type (Fig. 4E). The numbers of antral follicles appeared to be inversely correlated with *Nr5a1* expression levels, with the Tg-S ovaries containing fewer antral follicles than the Tg-A ovaries (Fig. 4E). To determine whether this was caused by reduced proliferation or increased apoptosis of follicular granulosa cells, we analysed the expression of Ki-67, a maker of proliferating cells, and cleaved Caspase 3, a marker of apoptotic cells, in the transgenic ovaries at P28. We observed no obvious changes in the follicles containing Ki-67-positive granulosa cells (Fig. 4J-L), but a significant increase in secondary follicles (type 4 and 5) containing cleaved Caspase 3-positive granulosa cells in both Tg-A and Tg-S ovaries (Fig. 4F, G–I, M–O). The increase in follicular atresia correlated with *Nr5a1* expression levels, with Tg-S ovaries exhibiting higher levels of atresia than Tg-A ovaries (Fig. 4F,G-I,M-O). No increase in type 6 antral follicles containing cleaved Caspase 3-positive cells was observed (Fig. 4F). Together, these results suggest that overexpression of *Nr5a1* caused increased apoptosis of granulosa cells in secondary follicles in a dose-dependent manner, resulting in reduced numbers of antral follicles.

**Figure 4:**
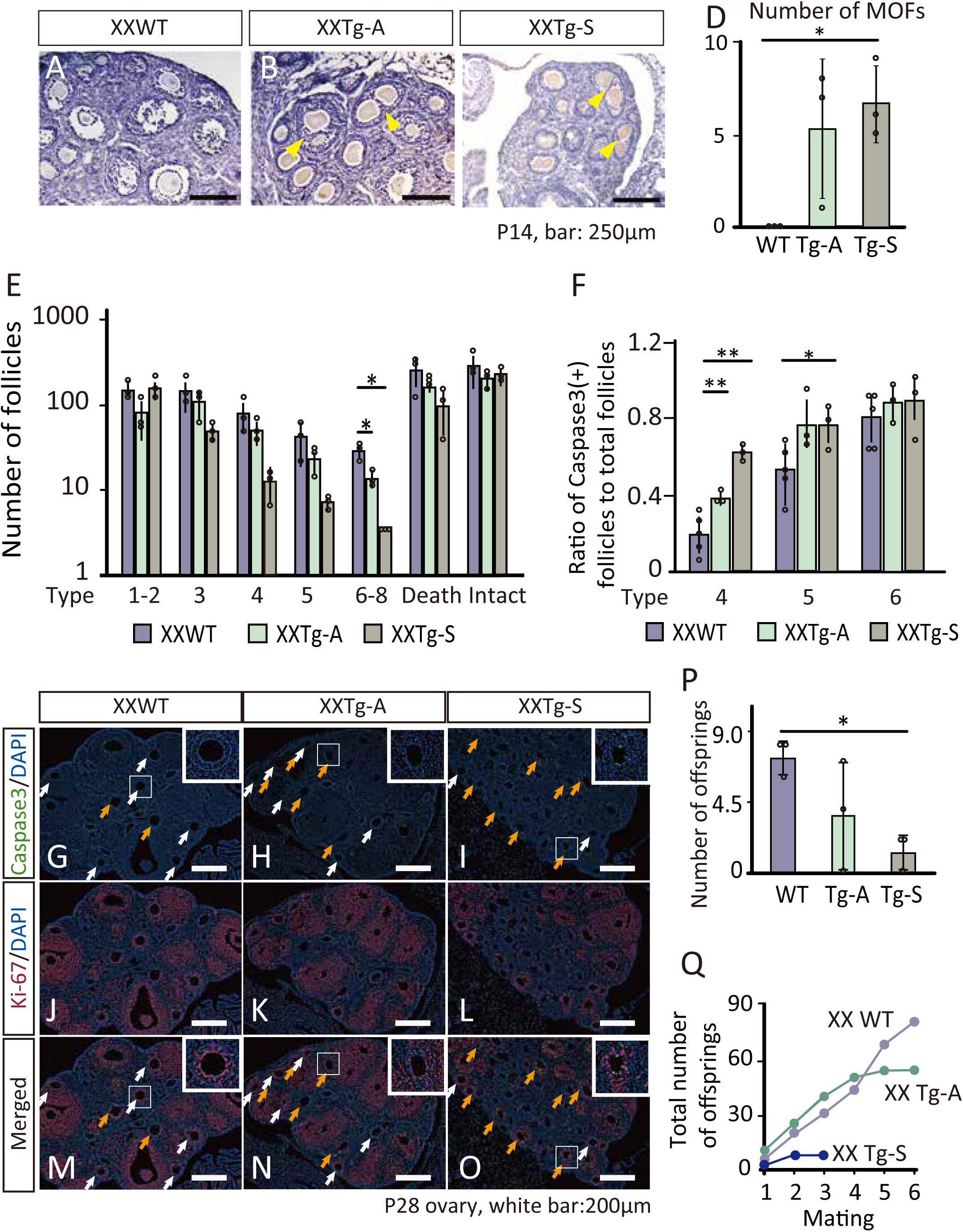
Impaired ovarian development in *Nr5a1* transgenic mice. (A–C) Histological analysis of ovaries in the wild-type, Tg-A, and Tg-S females. Multi-oocyte follicles (MOFs) were observed in Tg-A and Tg-S ovaries (arrowhead). (D) Numbers of MOFs were counted from serial ovarian sections. Mean ± SEM, n = 3. \**p* < 0.05 (Welch‘s test). (E) The numbers of antral follicles (type 6–8) significantly decreased in transgenic mice. The numbers of follicles of each type [1-2 (primordial), 3 (primary), 4-5 (secondary), 6-8 (antral), death and intact] were counted from serial ovarian sections. Mean ± SD, n = 3. \**p* < 0.05 (Welch‘s *t*-test). (F) Quantitation of secondary follicles containing cleaved Caspase 3-positive cells (G-I). The percentage of cleaved Caspase 3-positive follicles was plotted. Mean ± SEM, n = 3. \**p* < 0.05, \*\**p* < 0.01 (Welch‘s *t*-test). (G–O) Immunofluorescence analysis for markers of cell proliferation (Ki-67: red) or apoptosis (cleaved Caspase 3: green) in the XX gonads of the wild-type, Tg-A and Tg-S mice. Yellow and white arrows indicate secondary follicles (type 4–5) with or without cleaved Caspase 3-positive cells, respectively. (P) Average litter size of the first mating of wild-type, Tg-A, and Tg-S female mice (n = 3). Mean ± SEM. \**p* < 0.05 (Welch‘s *t*-test). (Q) Total number of progeny from wild-type, Tg-S and Tg-A female mice (n = 3).

We next assessed the fertility of Tg-A and Tg-S female mice. Tg-S female mice were almost infertile. Of three Tg-S females analyzed, the litter size was reduced from their first mating (Fig. 4P). The reproductive performance of Tg-A females was also reduced. Although Tg-A mice produced four consecutive litters to begin the study, they failed to produce additional litters thereafter (Fig. 4Q), suggesting that these females underwent premature ovarian insufficiency.

### *Nr5a1* overexpression represses Notch signaling levels in fetal ovaries

Interactions between germ cells and somatic pregranulosa cells are crucial for the formation of ovarian follicles, and Notch signaling plays a major role in mediating this interaction [29-31]. In the developing ovaries, oocytes and other neighbouring cells secrete the Notch ligands, including JAG1 and possibly JAG2, which bind to Notch receptors (mainly NOTCH2) present on the surface of pre-granulosa cells, thereby activating Notch signaling [32]. As a result, pre-granulosa cells proliferate and encapsulate individual germ cells to form primordial follicles, i.e., the resolution of germ cell syncytia [30]. Genetic ablation of either *Jag1* or *Notch2*, two important Notch pathway components, gives rise to MOFs and causes premature reproductive senescence [29]. Because of the similarity in the phenotypes between our *Nr5a1* transgenic mice and *Jag1*-or *Notch2*-deficient mice, we hypothesized that enforced expression of *Nr5a1* may compromise ovarian development by repressing the Notch signaling pathway.

We therefore analysed mRNA expression levels of a number of genes involved in Notch signaling in 15.5-dpc fetal ovaries using qRT-PCR (Fig. 5 A∼H). Supporting our hypothesis, we found significant down-regulation of several Notch pathway genes, including *Notch2, Notch3* and *Dll4*, in Tg-A ovaries compared with wild-type ovaries (Fig. 5B, C, H).

**Figure 5:**
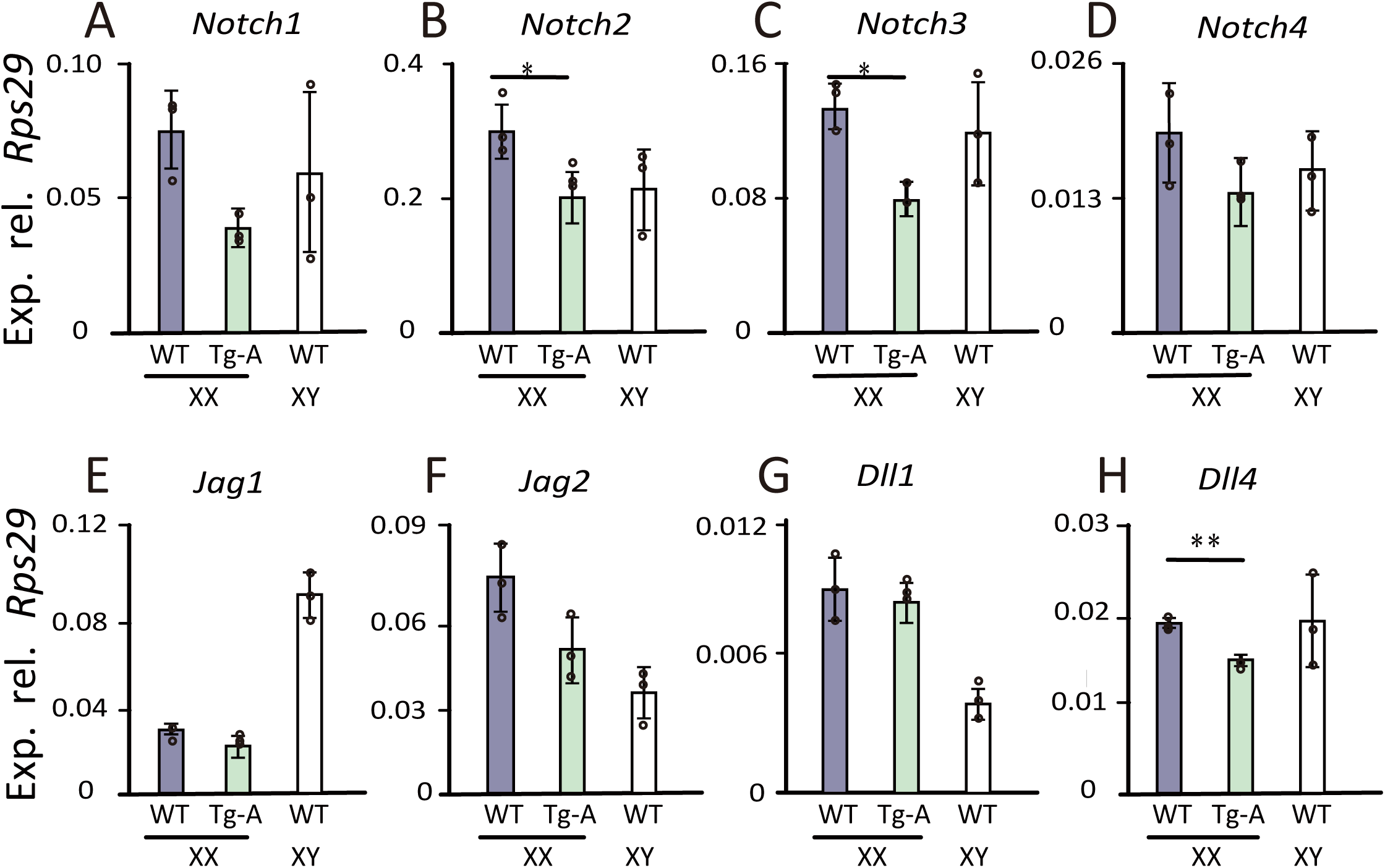
Expression of the Notch signaling pathway genes in XX Tg-A mice. (A-H) qRT-PCR analysis was performed on total RNA extracted from fetal ovaries of wild-type, Tg-A, and Tg-S mice at 15.5 dpc. Expression levels were normalized to *Rps29*. Mean ± SEM, n = 3. \**p* < 0.05, \*\**p* < 0.01 (Welch‘s *t*-test).

### A Notch2 agonist rescues the ovarian phenotype in Tg-A mice

To further clarify the contribution of Notch signaling to the ovarian phenotype in *Nr5a1* transgenic mice, we attempted a rescue experiment with a Notch2 agonist HMN2-29, a hamster monoclonal antibody [33]. We injected three doses of HMN2-29 or control hamster IgG intraperitoneally into pregnant mice carrying wild-type or Tg-A fetuses at 13.5, 16.5 and 18.5 dpc (Fig. S2A), and analysed the ovaries postnatally (P14 and P28). No obvious adverse effects of *in utero* administration of the Notch2 agonist were observed, as the treated mice appeared grossly normal with body weight comparable to wild type at P28 (Fig. S2B).

Confirming our hypothesis, we found reduced number of MOFs in Tg-A mice at P14 upon HMN2-29 treatment (Fig. 6A). Moreover, treatment with the NOTCH2 agonist completely rescued the numbers of antral follicles in the Tg-A ovaries at P28 (Fig. 6B). The complete rescue of antral follicle numbers appeared to be a result of reduced atresia of secondary follicles in the treated ovaries (Fig. 6C-O). These results suggest that *Nr5a1* fine-tunes Notch signaling levels in fetal ovaries to ensure proper folliculogenesis and normal fertility, and that repression of *Nr5a1* during fetal ovarian development is essential to allow the Notch signaling levels to elevate appropriately.

**Figure 6:**
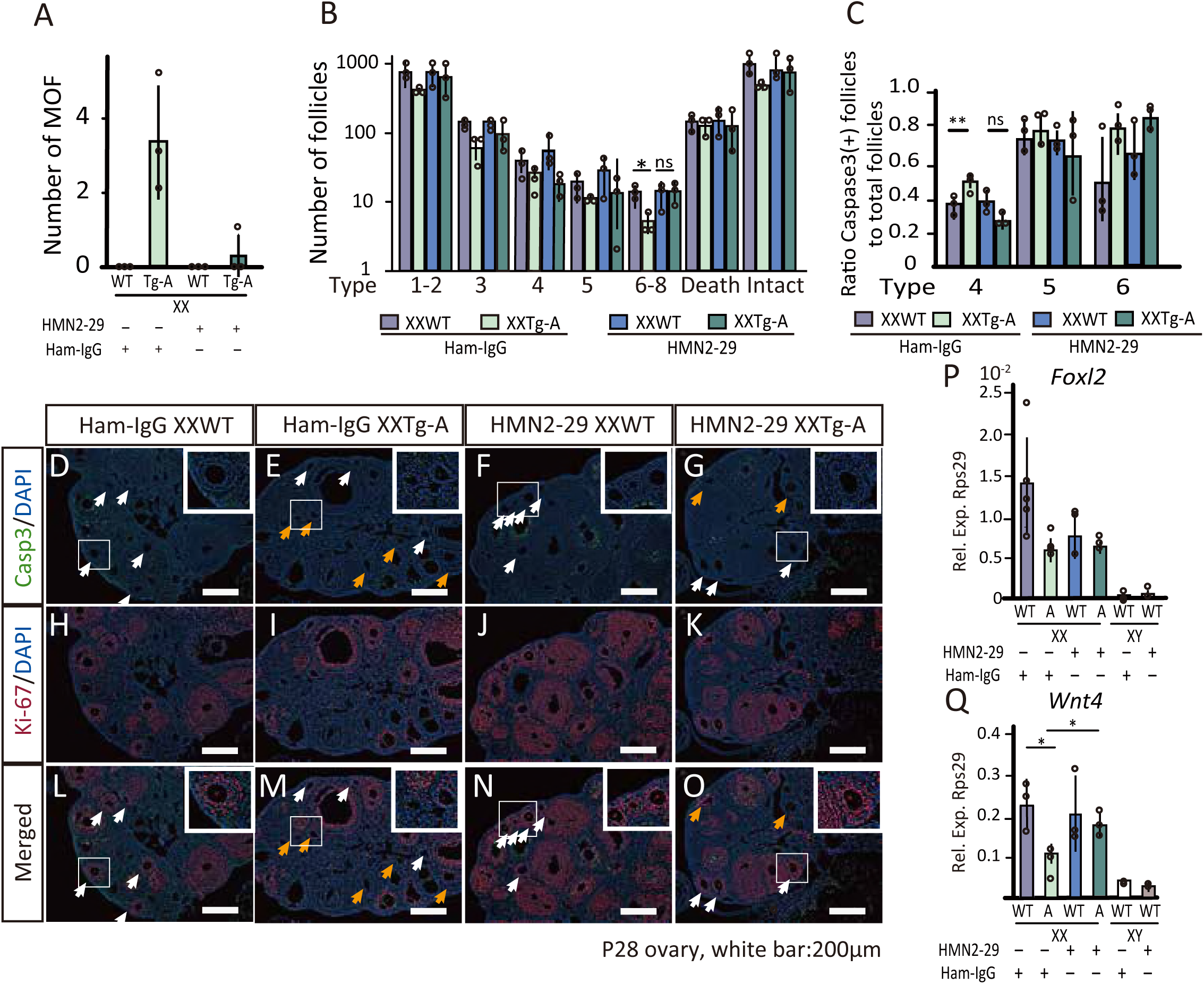
Administration of a Notch2 agonist rescued the ovarian phenotype in Tg-A mice. (A,B) The Notch2 agonist HMN2-29 or control hamster IgG was *in utero* administered to wild-type or Tg-A mice, and the ovarian phenotype was analyzed at P14 (A) and P28 (B). HMN2-29 administration almost completely rescued the formation of MOFs in the Tg-A ovaries at P14 (A), and restored the antral follicles to the wild-type level at P28 (B). Mean ± SEM, n = 3. \**p* < 0.05, \*\**p* < 0.01 (Welch‘s *t*-test). (C-O) Immunofluorescence analysis at P28 for markers of cell proliferation (Ki-67: red) or apoptosis (cleaved Caspase 3: green) in XX gonads of wild-type and Tg-A mice treated with hamster IgG or HMN2-29. White and yellow arrows indicate secondary follicles (type 4–5) with or without cleaved cleaved Caspase 3-positive granulosa cells, respectively. Quantitation of Caspase 3-positive follicles in (D-G) is shown in (C). Mean ± SEM, n = 3. \*\**p* < 0.01 (Welch‘s *t*-test); ns, not significant. (P,Q) qRT-PCR analysis for *Foxl2* (P) and *Wnt4* (Q) in P0 gonads of wild-type or Tg-A mice treated with hamster IgG or HMN2-29, respectively. Expression levels were normalized to *Rps29*. Mean ± SEM, n = 3. \**p* < 0.05, \*\**p* < 0.01 (Welch‘s *t*-test); ns, not significant.

We further explored molecular mechanisms by which the Notch signaling pathway regulates fetal ovarian development. To this end, we analysed expression levels of two master regulators of ovarian development, *Wnt4* and *Foxl2*, in the Tg-A ovaries at P0. We found that *Wnt4*, but not *Foxl2*, was significantly down-regulated in Tg-A mice compared to wild type (Fig. 6P, Q). Importantly, *Wnt4* expression in the Tg-A ovaries were fully restored by the administration of HMN2-29 (Fig. 6Q), suggesting that down-regulated *Wnt4* expression was a result of de-regulated Notch signaling pathway rather than a direct effect of NR5A1 overexpression. We note that this regulatory relationship is likely limited to the late-stage fetal ovaries, as expression of the Notch pathway genes starts in fetal ovaries from about 15.5 dpc [29].

## DISCUSSION

*Nr5a1* plays known essential roles in the development of genital ridges in both sexes and fetal testes in the male. We show in the present study that it also plays an important role in fetal ovarian development by fine-tuning the levels of Notch signaling.

### Overexpression of *Nr5a1* alone is insufficient to drive testis determination and differentiation in mice

Of the two transgenic lines generated, the Tg-S line, expressed *Nr5a1* in the XX gonads at levels comparable to that in wild-type testes. Nevertheless, XX mice of the Tg-S line developed female internal and external genitalia. We presume that NR5A1 may play different roles with respect to sex determination in humans and mice, and wild-type NR5A1 may not possess the ability to drive testis differentiation on its own in mice. Consistently, XX mice carrying heterozygous or homozygous R92W mutation in *Nr5a1* showed no signs of masculinization [13], supporting that NR5A1 (and the R92W mutant) may function differently in humans and mice. The differences between humans and mice regarding the molecular mechanisms of sex determination and gonadal development have been documented in several cases. For example, duplication of *DAX-1* (*NR0B1*), an orphan nuclear receptor gene, causes XY female sex reversal in humans [34], whereas transgenic overexpression of *Dax1* in mice does not normally cause female sex reversal [17]. Since a major function of DAX1 is to antagonize NR5A1 [1, 34-36], it is conceivable that NR5A1 also plays a species-specific role in sex determination.

### NR5A1 promotes fetal Leydig cell differentiation by restricting Notch signaling

Our results reveal a novel function of NR5A1 in the negative regulation of the Notch signaling pathway during fetal ovarian development. In addition to its critical functions in ovarian development, Notch signaling also plays a pivotal role in fetal testis development, particularly in the differentiation of fetal Leydig cells. Notch signaling restricts fetal Leydig cell differentiation by promoting and maintaining the progenitor cell fate [37]. Interestingly, NR5A1 has been suggested to promote fetal Leydig cell differentiation by suppressing Notch signaling in this context [38].

Consistent with these reports, we found that overexpression of NR5A1 in the fetal ovaries of Tg-S mice led to the differentiation of 3β-HSD positive cells, presumably ectopic fetal Leydig cells, at 15.5 dpc. However, the presence of these cells did not lead to an increase in plasma testosterone levels at birth. There may be too few 3β-HSD positive cells in the transgenic ovaries: this is borne out by the fact we did not find significant elevation of *Hsd3b* gene expression in the Tg-S fetal ovaries by qRT-PCR analysis (data not shown). A recent report revealed that fetal Leydig cells do not have the capacity to produce testosterone because they do not express 17β-hydroxysteroid dehydrogenases (17β-HSD), essential for the last step of testosterone synthesis [24, 39]. Hence, although some ectopic 3β-HSD-positive presumptive fetal Leydig cell differentiation occurred in the Tg-S mice, the absence of Sertoli cells expressing 17β-HSD (which convert androstenedione to testosterone) means that no testosterone can be produced.

### Fetal gonadal development requires optimal levels of *Nr5a1* and Notch signaling

Based on our and published data, we propose a model of NR5A1 function in fetal gonadal development (Fig. 7). The sexually dimorphic expression pattern of *Nr5a1* in the developing fetal gonads allows the Notch signaling activity to be tuned to optimal levels to suit distinct developmental programs. In fetal ovaries, downregulated *Nr5a1* de-represses Notch signaling, thereby allowing appropriate follicular development. On the other hand, elevated levels of NR5A1 in fetal testes represses Notch signaling, allowing fetal Leydig cells to differentiate. In ovarian development, the regulation of Notch signaling by NR5A1 may be context-dependent and limited to the fetal stage, since the mRNA levels of *Notch2/3* and *Nr5a1* are known to simultaneously increase after birth [29, 40].

**Figure 7:**
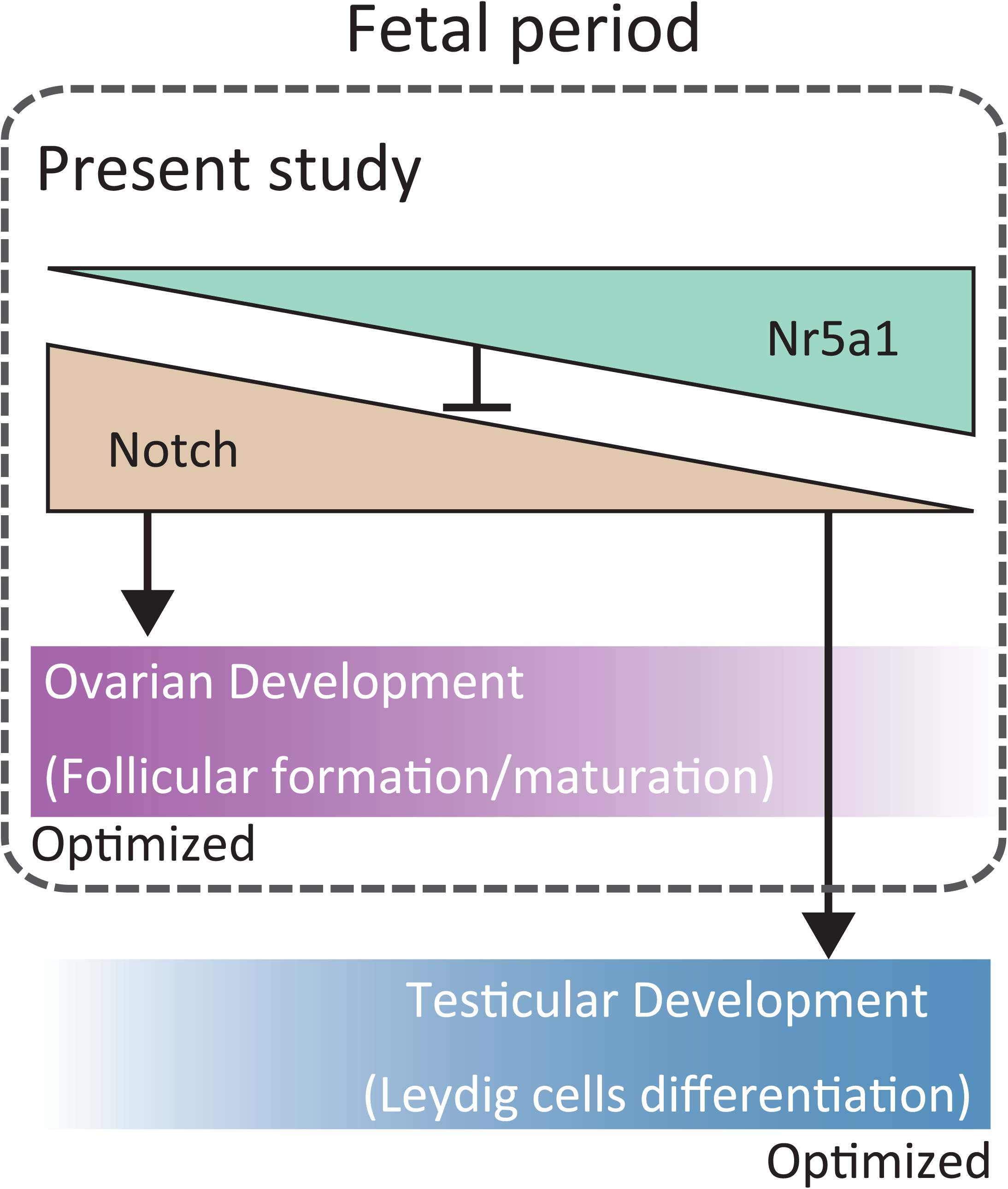
A model for NR5A1 function in fetal gonadal development in mice. NR5A1 fine-tunes Notch signaling levels to achieve optimal developmental outcomes in both fetal ovaries and testes.

Our model suggests that an optimal level of *Nr5a1* in fetal ovaries is required for proper development of follicles, and that either insufficient or excessive *Nr5a1* expression in fetal ovaries leads to impaired ovarian development. Consistent with this model, *Cited2*-null mice with severely reduced *Nr5a1* expression in fetal ovaries showed impaired expression of several ovarian marker genes [41], suggesting that low levels of *Nr5a1* expression are required for ovarian development. Similarly, women carrying loss-of-function mutation in *NR5A1* often develop premature ovarian insufficiency [42]. On the other hand, mild upregulation of *Nr5a1* expression in Tg-A mice was sufficient to impair follicular development and female fertility, even though the increase in expression was very mild compared to wild-type ovaries.

In summary, our study provides novel insight into the molecular pathways regulating fetal ovarian development, about which little is currently understood. We show that insufficiently repressed *Nr5a1* during fetal ovarian development leads to compromised follicular development and fertility issues due to dysregulated Notch signaling. Alterations in *Nr5a1* dosage, either reduced or excessive, result in pathological effects in ovarian development and female fertility, indicating that the precise control of *Nr5a1* at the transcriptional level is essential for optimal ovarian development. Further studies are required to reveal molecular details of the NR5A1-Notch-WNT4 axis in fetal ovarian development. We envisage that the improved understanding of how this pathway regulates ovarian development and female fertility would aid the development of artificial somatic ovarian cells, which in turn may provide a valuable treatment option in reproductive medicine.

## MATERIALS AND METHODS

### Generating transgenic mice

The mouse transgenesis procedure was based on a protocol described previously [18]. Briefly, a sequence containing the mouse *Nr5a1* coding region followed by an internal ribosomal entry site and the sequence encoding enhanced green fluorescent protein (*Nr5a1*-IRES-*Egfp*) was cloned into the PBWt1-Dest vector via Gateway LR recombination (Fig. 1A). Transgenic founder mice were produced by pronuclear injection of the PBWt1-*Nr5a1*-IRES-*Egfp* vector DNA and hyperactive piggyBac transposase mRNA as described. The XY male founders mated with BDF1 females, and transmitted the transgene through the germ line. For subsequent analyses, we chose two lines, Tg-A and Tg-S. Both lines were maintained by mating F0 or F1 XY transgenic male mice with BDF1 wild type females. Genotyping and sexing was performed by PCR (primer sequences provided in Supplementary Table S1) using genomic DNA prepared from tail or ear biopsies. All animal procedures were approved by the Institutional Animal Care and Use Committee of Tokyo Medical and Dental University.

### Fertility analysis of transgenic female mice

The fertility of Tg-A and Tg-S female mice, compared with wild type female mice, was assessed after they turned 6 weeks old by continuous mating with approximately 30-week old BDF1 male mice. Three female mice from each line were analyzed. The average number of offspring in the first litter and the aggregated number of offspring from all litters were calculated for each genotype.

### Real-time RT-PCR analysis

RNA was extracted from 15.5 dpc or P0 mice gonads using QIAGEN RNeasy Mini kit according to manufacturer‘s instructions. Typically, 0.2 µg of total RNA was reverse transcribed using random hexamers (Promega) and SuperScript III reverse transcriptase (Invitrogen). Quantitative real-time PCR (qRT-PCR) was conducted using a Light Cycler system (Roche Diagnostics, Basel, Switzerland) with the Light Cycler DNA master SYBR Green kit (Roche) for 45 cycles. Gene expression levels were analysed using the comparative cycle time (Ct) method. Primers used in these experiments are listed in Supplementary Table S1. *Rps29* (Ribosomal protein S29) served as the housekeeping gene for normalization, as it shows minimum variability during fetal gonadal development [36].

### Histological analyses

#### Cryosections

Gonadal samples were fixed overnight in 4% paraformaldehyde (PFA) at 4 °C. After washing three times with PBS, samples were incubated overnight in 20% sucrose/PBS at 4 °C. Samples were then incubated in 3/4 30% sucrose/OCT at 40 °C for 30 min and embedded.

#### Paraffin-sections

Gonadal samples were fixed overnight in 4% PFA at 4 °C and embedded in paraffin. The blocks were sectioned at 6-µm thickness and were later deparaffinised as previously described [43].

#### Hematoxylin and Eosin (HE) stain

After staining with 2× Haematoxylin for 4 min, the sections were washed for 12 min under running water and then stained with 1.0% Eosin for 2 min.

#### Immunofluorescence (IF)

We employed cryosections and paraffin sections for IF. For cryo-sections, 8 µm of samples were washed twice with PBS and activated with citric acid solution. The sections were blocked with 5% BSA-PBS at room temperature for 1 h, followed by overnight incubation with primary antibody at 4 °C. Next, the sections were washed twice with PBS-T, incubated with secondary antibody at room temperature for 1.5 h, and again washed twice with PBS-T. Finally, the sections were incubated with DAPI (1:1000; Dojindo) for 5 min, washed once with PBS-T, and mounted in Fluoromount™. The information of antibodies used in this study was indicated in Supplementary Table S2.

#### Counting numbers of follicles, apoptotic follicles, or multi-oocyte follicles (MOFs)

Counting was performed as previously described [31]. Briefly, serial sections (6-µm thick) from a whole ovary were placed on 5 slides, each slide containing sequential slices at every 30 µm interval (6 µm * 5 slides). Multi-oocytic follicles were defined as follicles containing more than a single oocyte. Anti-MVH antibody (ab13840, Abcam) and DAB staining (25985-50, Nacalai Tesque Inc.) was used to count the number of follicles and MOFs. The types of follicles were classified according to previous reports [44, 45]. We used anti-Cleaved Caspase 3 (Cell signaling technology, Cat No. 9579) to identify apoptotic granulosa cells and calculated the percentage of Caspase 3-positive follicles.

### Hormone measurements

Free testosterone and estradiol levels of P0 mice were measured using commercially available ELISA kits (IBL International (DB52181) and Cayman Chemical (582251)). Three sets of plasma samples obtained from each of 5 mice were analyzed according to the manufacturer‘s protocol.

### Preparation and administration of the Notch agonist HMN2-29

BDF1 eggs were fertilized *in vitro* with Tg-A sperm and transplanted into the oviducts of ICR mice. The Notch2 agonist HMN2-29, a hamster monoclonal antibody, was prepared as described previously [33]. Three doses of 0.5 mg of HMN2-29 or control hamster IgG (Wako, Cat No. 145-19561) were intraperitoneally injected into pregnant ICR mice at 13.5, 16.5 and 18.5 dpc.

### Statistical analysis

We used the unpaired *t*-test (Welch‘s test) to determine statistically significant differences between the test and the control group.

## Data availability

Authors can confirm that all relevant data are included in the paper and/ or its supplementary information files

## Acknowledgements

This study was partially supported by the grant, KAKENHI, Kiban (C) (15K09615) funded by Japan Society for the Promotion of Science (JSPS), for K.K. The funders had no role in study design, data collection and analysis, decision to publish, or preparation of the manuscript.

## Author Contributions

KK, RN, HS, YK, MKA, TM contributed to the conception and the design of the present study. Acquisition of data, analysis and interpretation of data were performed by RN, HS and KK. The manuscript was drafted by KK, TM and RN, and critically revised by LZ, JB and PK. HS, LZ, JB, PK, MT, MKA and HY provided the materials for the present study, including Piggy BAC construct and Notch agonist. All experiments were performed by RN, HS, LZ, KK and AHT. All authors approved the final version to be published

---

**Supplementary Figure S1: Overexpression of *Nr5a1* affected the morphology of fetal ovaries in the Tg-A mice.** (A,B) Bright-field images of 15.5 dpc-ovaries from Tg-A and wild-type mice. (C,D) Scatter plot measuring the length or width of the fetal ovaries. \**p* < 0.05 (Welch‘s *t*-test). (E) qRT-PCR analysis showing that *Sox4* was significantly down-regulated in Tg-A fetal ovaries compared with wild type ovaries at 15.5 dpc. Expression levels were normalized to *Rps29*. Mean ± SEM, n = 3. \**p* < 0.05 (Welch‘s *t*-test).

**Supplementary Figure S2: *In utero* administration of the Notch2 agonist did not affect body weight.** (A) Schematics showing the experimental design. The Notch2 agonist HMN2-29 was administered *in utero* at 13.5, 16.5 and 18.5 dpc via i.p. injection. Hamster IgG was used as the mock control. (B) *In utero* treatment of HMN2-29 did not cause a significant change in mouse body weight at P28. ns, not significant (Welch‘s *t*-test).

**Supplementary Table S1.** The list of primer sets used in this study

**Supplementary Table S2.** The details of antibodies used in this study

